# A 3-D Constitutive Model for Finite Element Analyses of Agarose with a Range of Gel Concentrations

**DOI:** 10.1101/2020.07.25.221317

**Authors:** Xiaogang Wang, Ronald K. June, David M. Pierce

## Abstract

Hydrogels have seen widespread application across biomedical sciences and there is considerable interest in using hydrogels, including agarose, for creating *in vitro* three-dimensional environments to grow cells and study mechanobiology and mechanotransduction. Recent advances in the preparation of agarose gels enable successful encapsulation of viable cells at gel concentrations as high as 5%. Agarose with a range of gel concentrations can thus serve as an experimental model mimicking changes in the 3-D microenvironment of cells during disease progression and can facilitate experiments aimed at probing the corresponding mechanobiology, e.g. the evolving mechanobiology of chondrocytes during the progression of osteoarthritis. Importantly, whether stresses (forces) or strains (displacement) drive mechanobiology and mechanotransduction is currently unknown. We can use experiments to quantify mechanical properties of hydrogels, and imaging to estimate microstructure and even strains; however, only computational models can estimate intra-gel stresses in cell-seeded agarose constructs because the required *in vitro* experiments are currently impossible. Finite element modeling is well-established for (computational) mechanical analyses, but accurate constitutive models for modeling the 3-D mechanical environments of cells within high-stiffness agarose are currently unavailable. In this study we aimed to establish a 3-D constitutive model of high-stiffness agarose with a range of gel concentrations. We applied a multi-step, physics-based optimization approach to separately fit subsets of model parameters and help achieve robust convergence. Our constitutive model, fitted to experimental data on progressive stress-relaxations, was able to predict reaction forces determined from independent experiments on cyclical loading. Our model has broad applications in finite element modeling aimed at interpreting mechanical experiments on agarose specimens seeded with cells, particularly in predicting distributions of intra-gel stresses. Our model and fitted parameters enable more accurate finite element simulations of high-stiffness agarose constructs, and thus better understanding of experiments aimed at mechanobiology, mechanotransduction, or other applications in tissue engineering.

## 1. Introduction

Hydrogels, consisting of hydrophilic polymers in aqueous environments, are relatively easy to work with and have seen widespread application across biomedical sciences, e.g. in delivering drugs, and developing organoids and organs-on-a-chip (Elviri et al., 2017; Zhang and Khademhosseini, 2017; Liu et al., 2019). Recent technical advances generated hydrogels that exhibit both greater stiffness and toughness to further develop the application space, particularly in areas previously limited by low mechanical stiffness (Xu et al., 2019). There is considerable interest in using hydrogels, including agarose, for creating *in vitro* three-dimensional environments to grow cells and study mechanobiology, mechanotransduction, or applications in tissue engineering (Hung et al., 2004; Mauck et al., 2007). Studies leveraging agarose to study cells and mechanotransduction used concentrations up to 5% weight per volume where the stiffness of agarose increases with concentration (Zignego et al., 2014; Jutila et al., 2015).

Cartilage lines the bony interfaces within articulating joints to facilitate load transfer and smooth joint movement. In osteoarthritis (OA), the most common degenerative joint disease, cartilage weakens and deteriorates, leading to pain, disability, and eventually total joint replacement. Chondrocytes, the only cells within articular cartilage, reside within a gel-like pericellular matrix (PCM) within the extracellular matrix. PCM serves to transmit joint-level loads to the embedded chondrocytes and also presents progressively reduced stiffness with advancing OA.

Recent advances in the preparation of agarose gels enable successful encapsulation of viable cells at gel concentrations as high as 5%. These agarose specimens also present equilibrium moduli within the range of those measured for healthy PCM surrounding chondrocytes *in vivo*, on the order of 42 kPa (Jutila et al., 2015). In fact, changes in the stiffnesses of agarose microenvironments generate corresponding changes in mechanotransduction of the embedded chondrocytes (Alexopoulos et al., 2005; Darling et al., 2010). Since the stiffness of the PCM decreases with OA, agarose with a range of gel concentrations can serve as an experimental model mimicking osteoarthritic changes in the 3-D microenvironment of the PCM in cartilage and facilitate experiments aimed at probing the mechanobiology of OA (McCutchen et al., 2017). We recently quantified the relationship between concentration and stiffness for agarose, and thus established an *ex vivo* experimental model mimicking both the healthy and osteoarthritic PCM (Jutila et al., 2015).

Importantly, whether stresses (forces) or strains (displacement) drive mechanotransduction is currently unknown. We can use experiments to quantify mechanical properties of hydrogels, and imaging to estimate microstructure and even strains, e.g. Chan et al. (2016); however, only computational models can estimate intra-gel stresses in cell-seeded agarose constructs because the required *in vitro* experiments are currently impossible. Finite element (FE) modeling is well-established for (computational) mechanical analyses, but accurate constitutive models for modeling the 3-D mechanical environments of cells within high-stiffness agarose are currently unavailable. In this study we aimed to establish a 3-D constitutive model of high-stiffness agarose with a range of gel concentrations. Our model and fitted parameters enable more accurate FE simulations of high-stiffness agarose constructs, and thus better understanding of experiments aimed at mechanobiology, mechanotransduction, or other applications in tissue engineering.

## 2. Materials and Methods

### 2.1. Experimental Evidence

We previously completed mechanical testing of agarose which can encapsulate cells and facilitate mechanobiological experiments (Jutila et al., 2015). Briefly, we prepared cylindrical constructs using low-gelling-temperature agarose (Sigma, Type VII-A A0701) for mechanical tests. We dissolved 3-5% (weight per volume) agarose in phosphate-buffered saline (PBS) at 1.1× strength and 40°C. After approximately five minutes we diluted the dissolved agarose to 1 × with PBS at 40°C. We then cast agarose using anodized aluminum molds at 23°C to produce cylindrical constructs of 3.0, 3.5, 4.0, 4.5, and 5% agarose (height: 12.7 ± 0.1 mm; diameter: 7.0 ± 0.1 mm). We completed mechanical testing using a custom-built bioreactor to apply displacement-controlled loads and established data for independent calibration and validation of our modeling efforts.

#### 2.1.1. Data for calibration

We first performed progressive stress-relaxation tests in unconfined compression. Prior to testing, we equilibrated constructs of 3.0 (*n* = 5), 3.5 (6), 4.0 (6), 4.5 (8), and 5% (7) agarose (*n* = 32 total constructs) in PBS at 37°C for 30 min. We then applied unconfined, uniaxial compression along the main axis of the constructs using three consecutive steps of 4% nominal compressive strain, and we held each strain step for 90 min at 37°C in tissue culture conditions (humidified with 5% CO_2_) for stress relaxation. We sampled time, displacement, and force at 1000 Hz for the duration of each test.

#### 2.1.2. Data for validation

We also performed separate cyclic, unconfined compression tests on an independent set of constructs. We first applied unconfined, uniaxial compression along the main axis of the constructs of 3.0 (*n* = 6), 3.5 (6), 4.0 (4), 4.5 (10), and 5% (6) agarose (*n* = 32 total constructs) at 5% prestrain for two hours. We then applied 100 cycles of sinusoidal compression from 3.1 to 6.9% nominal compressive strain at 0.55 Hz. We sampled time, displacement, and force at 100 Hz for the duration of each test to ensure sampling above the Nyquist limit.

### 2.2. Constitutive Modeling

We used the theory of porous media to describe agarose as a biphasic (poroelastic) continuum *φ* = *φ*^S^ + *φ*^F^ consisting of a solid phase *φ*^S^ (with an isotropic and statistically regular pore distribution) saturated with the fluid phase *φ*^F^. We formulated the total Cauchy stress tensor as (Pierce et al., 2016; Wang et al., 2018)

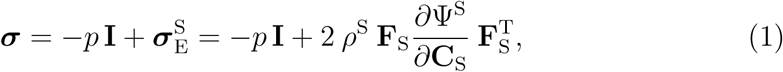

where *p* is the fluid pressure, **I** is the second-order identity tensor, 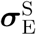 Cauchy stress tensor, *ρ*^S^ is the current partial density of the solid, **F**_S_ = ∂**x**_S_*/*∂**X**_S_ is the deformation gradient tensor of the solid, 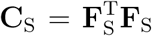 is the right Cauchy-Green tensor, and Ψ^S^ is the solid Helmholtz free-energy function. For generality we assumed a Mooney-Rivlin constitutive model for the solid phase such that (Simo and Pister, 1984)

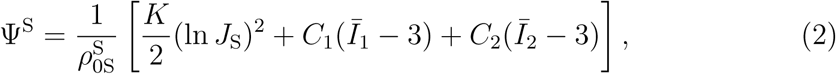

where 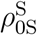 is the reference partial density of the solid, *K* [MPa] degenerates to a non-physical, penalty parameter used to enforce incompressibility, *J*_S_ = det**F**_S_ is the Jacobian determinant of the solid, *C*_1_ [MPa] and *C*_2_ [MPa] are model parameters with *µ* = 2(*C*_1_ + *C*_2_) corresponding to the shear modulus of the underlying matrix in the reference configuration, and *Ī*_1_ and *Ī*_2_ are the first and second invariants of the deviatoric right Cauchy-Green deformation tensor 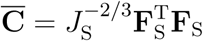.

We calculate the viscous contribution to the total stress in the Lagrangian configuration using the second Piola-Kirchhoff effective stress of the solid 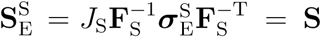. To determine the time-dependent (viscous) response of the solid we step through time *t* ∈ [0+, *T*] and focus on a representative subinterval [*t*_*n*_, *t*_*n*+1_] with Δ*t* = *t*_*n*+1_ − *t*_*n*_ the associated time increment (Holzapfel, 1996; Pierce et al., 2009). Algorithmically we know all the relevant kinematic quantities at times *t*_*n*_ and *t*_*n*+1_ and we can calculate the stress **S**_*n*_ at time *t*_*n*_ uniquely via (1)-(2). We then calculate the second Piola-Kirchhoff stress **S**_*n*+1_ at time *t*_*n*+1_ as

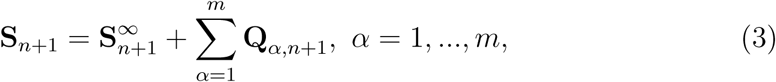

where 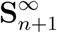 is the total elastic response computed from the given strain measures at *t*_*n*+1_ and **Q** _*α,n*+1_ are the non-equilibrium stresses associated with *α* = 1, …, *m*, viscoelastic (time-dependent) processes. We compute the non-equilibrium stresses assuming a linear evolution equation for each viscoelastic process

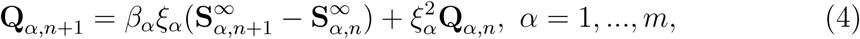

where *β*_*α*_ [-] is a dimensionless magnitude factor, *ξ*_*α*_ = exp(−Δ*t/*2*τ* _*α*_) with *τ* _*α*_ [s] the associated relaxation time, and **Q** _*α*,0_ = **0** for all *α* = 1, …, *m* (Holzapfel and Gasser, 2001).

To model the corresponding intrinsic permeability of the solid we assumed the isotropic Holmes-Mow permeability **K** such that (Holmes and Mow, 1990)

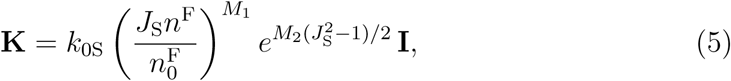

where *k*_0S_ [m^4^*/*(N *·* s)] is the reference hydraulic permeability, *n*^F^ and 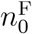 are the current and reference volume fractions of the fluid respectively, and *M*_1_ [-] and *M*_2_ [-] are dimensionless parameters controlling the deformation dependence of the permeability.

### 2.3. Model Selection and Calibration

We created a quarter-symmetry model of the cylindrical constructs using 20-node hexahedral elements in FEBio (University of Utah), Fig. 1(a)-(b). We finalized the mesh density using an *h*-refinement convergence study on the lateral displacements and axial Cauchy stresses, where the latter relates to the predicted total reaction forces. We applied symmetry boundary conditions to nodes on the cut faces of the quarter cylinder and fixed the *z*-displacement degrees of freedom of nodes on the bottom surface. We set the corresponding fluid flux normal to these surfaces to zero. We specified all remaining nodes in contact with the physiologic solution as free to displace and set the corresponding fluid pressure to zero (allowing fluid flux). We then prescribed the axial displacement of nodes on the top surface in the *z* direction to simulate unconfined compression.

**Figure 1:**
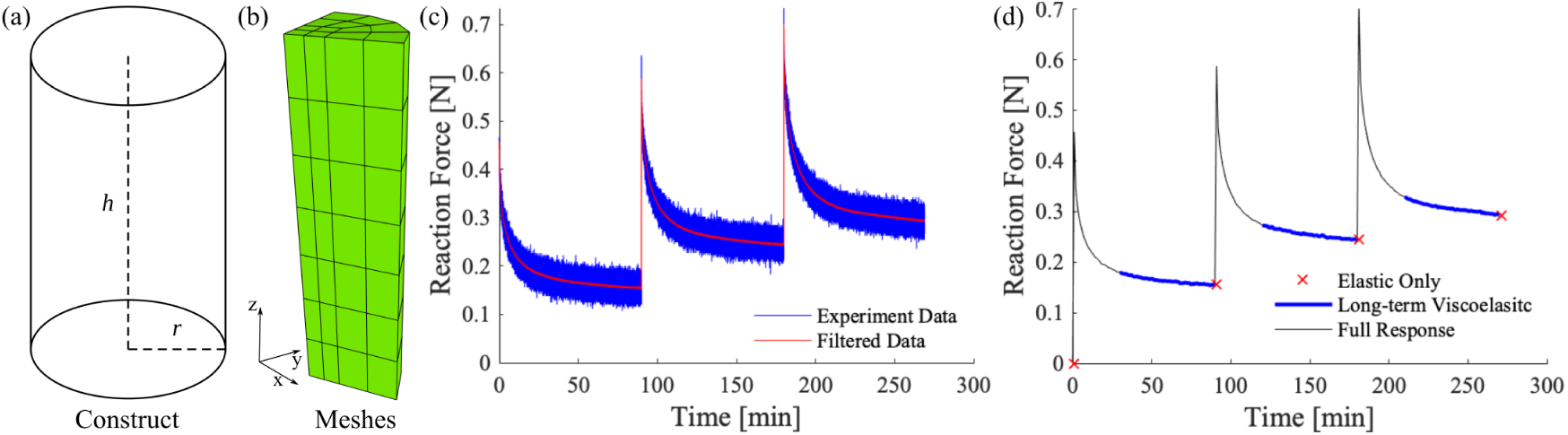
Finite element modeling of agarose constructs: (a) schematic image of the cylindrical constructs (*h* = 12.7 mm, *r* = 7.0 mm) and (b) the corresponding quarter-symmetry mesh. Representative data and elements of the parameter fitting: (c) raw and resampled data after smoothing with two moving-average filters and (d) separating mechanical responses within the full time-dependent data.

To prepare the mechanical data (Section 2.1) for the parameter fittings we filtered and resampled our data from the step-wise stress-relaxation tests. We first applied a moving-average filter with a window of length *b* = 10, and then applied a second moving-average filter with *b* = 20 to minimize noise. After resampling we obtained 90 data points for each load step which captured the peak reaction force under compression and the subsequent relaxation in one-minute increments thereafter. Our resampled data thus contained 271 points including one reference point before loading and 270 points encompassing the three progressive increments in strain, Fig. 1(c)-(d).

In light of the data available, and to minimize the computational burden, we determined the parameters related to permeability (*k*_0S_, *M*_1_, and *M*_2_, cf. (5)) prior to fitting the remaining parameters (NB. recall 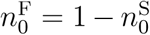). We calculated *k*_0S_ = *κ/µ*_v_ where *κ* [m^2^] is the Darcy permeability and *µ*_v_ [Pa·s] is the dynamic viscosity. We determined the Darcy permeability *κ* based on the agarose concentration 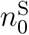 as (Johnson and Deen, 1996)

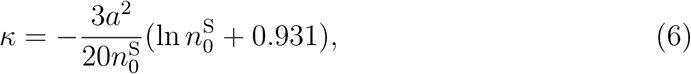

where *a* = 1.9 nm is the fiber radius (Djabourov et al., 1989) and 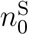 is the reference volume fraction of the solid (NB. not the same as *ϕ*, the concentration of the agarose gel). We also fixed *M*_1_ = 2 and *M*_2_ = 1.5.

We optimized the elastic and viscoelastic model parameters using the “Leven-berg-Marquardt” method within FEBio (Maas et al., 2012), see Appendix A. We first ran only elastic simulations to determine only the elastic parameters *C*_1_, *C*_2_.

Next, we fit the parameters related to the viscoelastic processes (VPs), using the fitted elastic parameters, with three different approaches:

1. we fitted all 271 data points in one step using one VP (*β*_1_, *τ*_1_) termed “One VP, One Step;”
2. we fitted all 271 data points in one step using a two VPs (*β*_1_, *τ*_1_ for a longer-term and *β*_2_, *τ*_2_ for a shorter-term process) termed “Two VPs, One Step;” and
3. we fitted the last 181 data points (two thirds of the total of each strain increment) for a longer-term VP (*β*_1_, *τ*_1_) and then fitted all 271 points by adding a shorter-term VP (*β*_2_, *τ*_2_) termed “Two VPs, Two Steps.”

To determine the best parameter set for each specimen (*n* = 32) we selected the fitting approach that generated the smallest objective functions *f*_*obj*_ [N^2^] for each of the 32 parameter optimizations. We presented the final parameter sets as means ± standard deviations for each concentration of agarose.

### 2.4. Model Validation

Leveraging our quarter-symmetry model of the cylindrical constructs, cf. Section 2.3, we changed only the boundary conditions applied to the top surface. Here we applied a 5% prestrain until equilibrium, and then applied a sinusoidal, cyclical displacement to generate axial, nominal strains ranging from 3.1 to 6.9%. For each specimen fitted in Section 2.3 (3.0, 3.5, 4.0, 4.5, and 5% agarose with *n* = 5 per concentration, 25 total simulations) we used the fitted model parameters and predicted the total reaction force both at equilibrium under 5% prestrain and as a function of time during cyclic loading. We simulated five cycles and compared the predicted reaction forces vs. time of the last three (repeatable) cycles against the experimental measurements (M ± SD) for each concentration. We validated our model predictions by comparing the shapes of the cyclical reaction forces (qualitatively), and by comparing the predicted reaction forces at 5% prestrain (*F*_0_) and both the peaks (*F*_P_) and valleys (*F*_V_) during cyclic loading (quantitatively).

### 2.5. Statistical Analyses

Prior to statistical analyses, we checked the data for normality using the Shapiro-Wilk test. To evaluate the performance of the models, i.e. which provided the best fits of the data, we used statistical analyses to compare between the various constitutive models and fitting procedures using the stress-relaxation data. We assessed fit by the sum of the squared error (the values of the objective functions) between the model predictions and the experimental data. After determining the constitutive model and fitting approach that resulted in the best fits to the experimental data we used a repeated measures ANOVA with Sidak’s multiple comparisons to assess the performance. To assess relationships between the fitted model parameters and the concentrations of agarose, we calculated linear correlation coefficients *r* for each parameter, which have positive values when the parameters increase with increasing agarose concentration. To quantify our independent validation of the fitted model, we also completed linear regressions (without a *y*-intercept) to quantify the predictive power of our simulation results (model predictions) against the experimental measurements using *R*-square values. We completed all statistical analyses using GraphPad Prism8 (La Jolla, CA, USA) and considered *p >* 0.05 as statistically significant.

## 3. Results

### 3.1. Model Selection and Calibration

We first determined the hydraulic permeability in the reference configuration *k*_0S_ as a function of the agarose concentration based on *κ* and *µ*_v_ (Johnson and Deen, 1996; Fernández et al., 2008), see Table 1.

**Table 1:** Permeability parameters of agarose gels with concentrations of 3-5%.

| $\phi[\%]$ | $\kappa[\text{mm}^2]$ | $\mu_v[\text{MPa} \cdot \text{s}]$ | $k_{0S}[\text{mm}^4/\text{N} \cdot \text{s}]$ |
| --- | --- | --- | --- |
| 3.0 | $4.65 \times 10^{-11}$ | $1.00 \times 10^{-6}$ | $4.65 \times 10^{-5}$ |
| 3.5 | $3.75 \times 10^{-11}$ | $1.75 \times 10^{-6}$ | $2.14 \times 10^{-5}$ |
| 4.0 | $3.10 \times 10^{-11}$ | $2.50 \times 10^{-6}$ | $1.24 \times 10^{-5}$ |
| 4.5 | $2.61 \times 10^{-11}$ | $3.25 \times 10^{-6}$ | $8.03 \times 10^{-6}$ |
| 5.0 | $2.24 \times 10^{-11}$ | $4.00 \times 10^{-6}$ | $5.59 \times 10^{-6}$ |

Next, we optimized the elastic and viscoelastic model parameters for all combinations of constitutive models and fitting approaches. The biphasic Mooney-Rivlin (not neo-Hookean, see Appendix B, Fig. 5) model with two viscoelastic processes fit using the one-step approach produced the best fits to the calibration data, i.e. the “Two VPs, One Step” approach, see Fig. 2.

**Figure 2:**
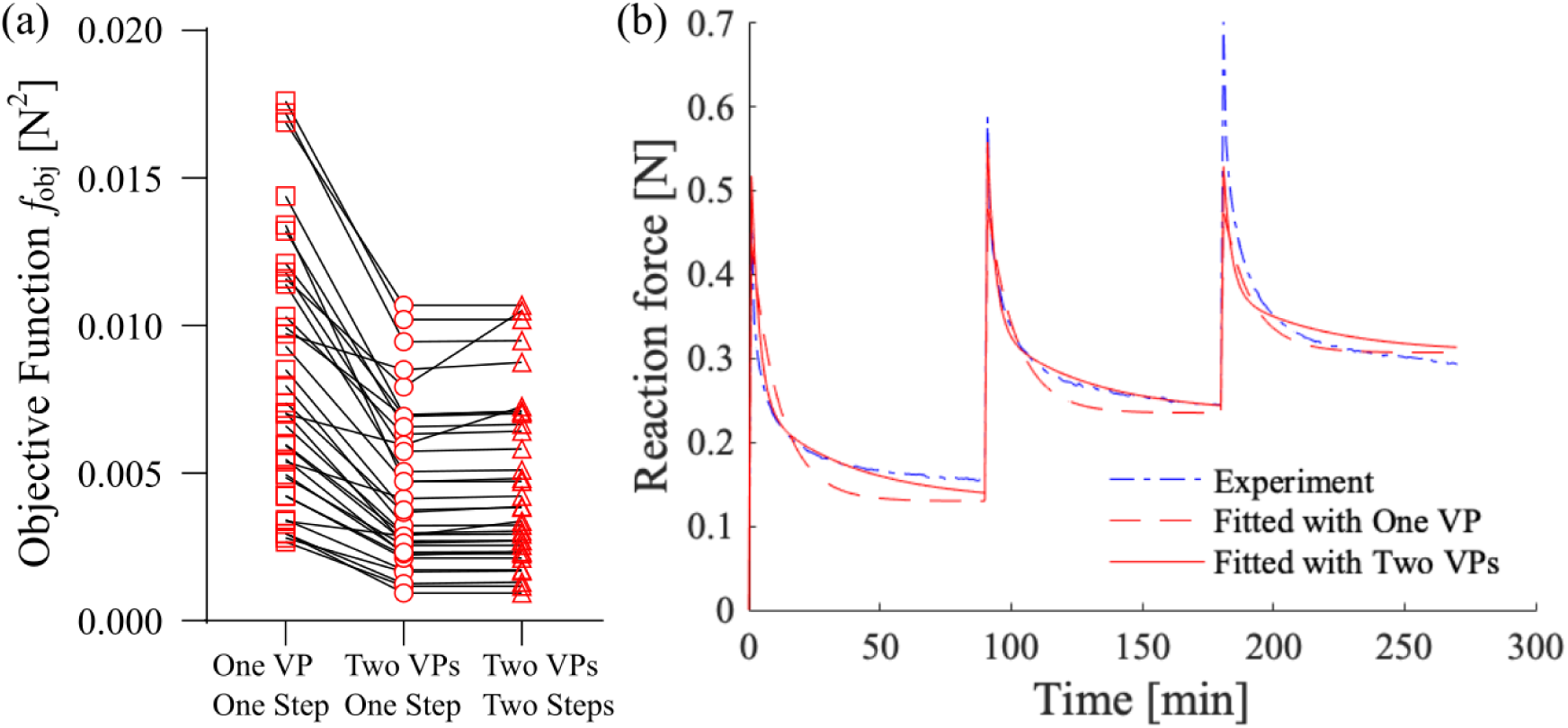
Direct comparison of the performance of the Mooney-Rivlin models including one or two viscoelastic processes and fitted with one- or two-step approaches: (a) values of the corresponding objective functions for all 32 specimens, (b) experimental data and predictions with fitted Mooney-Rivlin models (including one or two viscoelastic processes) for a representative agarose specimen.

More specifically, Fig 2(a) shows a direct comparison of the objective function values for each specimen fitted using specific models and approaches, while Fig 2(b) shows representative filtered experimental data and corresponding results from FE simulations applying the calibrated (optimized) parameters from a representative specimen. The model errors (i.e. objective functions) were lognormally distributed (all *p >* 0.13). Our repeated measures ANOVA identified a main effect (*p <* 0.0001), and *post hoc* multiple comparisons found that two VPs significantly improved the fitting quality versus that with only one VP (*p <* 0.00001), see Fig. 2(a,b). However, sequentially fitting the long-term and short-term VPs did not improve the fitting quality, see Fig. 2(a).

We presented the fitted parameters for the biphasic, viscoelastic Mooney-Rivlin model as a function of agarose concentration in Table 2, where the shear modulus *µ* = 2(*C*_1_ + *C*_2_).

**Table 2:** Elastic and viscoelastic parameters for the Mooney-Rivlin model of agarose gels with concentrations of 3-5%.

| $\phi$[%] | $C_1$[MPa] | $C_2$[MPa] | $\mu$[MPa] | $\beta_1$[-] | $\tau_1$[s] | $\beta_2$[-] | $\tau_2$[s] |
| --- | --- | --- | --- | --- | --- | --- | --- |
| 3.0 | 0.0284<br>$\pm 0.0023$ | -0.0208<br>$\pm 0.0018$ | 0.0152<br>$\pm 0.0011$ | 0.59<br>$\pm 0.17$ | 2566.07<br>$\pm 650.65$ | 3.09<br>$\pm 0.54$ | 126.55<br>$\pm 13.93$ |
| 3.5 | 0.0324<br>$\pm 0.0017$ | -0.0230<br>$\pm 0.0016$ | 0.0187<br>$\pm 0.0020$ | 0.75<br>$\pm 0.28$ | 2141.78<br>$\pm 352.07$ | 2.65<br>$\pm 0.62$ | 115.14<br>$\pm 27.86$ |
| 4.0 | 0.0368<br>$\pm 0.0024$ | -0.0256<br>$\pm 0.0019$ | 0.0223<br>$\pm 0.0014$ | 0.98<br>$\pm 0.28$ | 1865.01<br>$\pm 438.82$ | 2.51<br>$\pm 0.52$ | 149.02<br>$\pm 23.52$ |
| 4.5 | 0.0527<br>$\pm 0.0042$ | -0.0379<br>$\pm 0.0037$ | 0.0297<br>$\pm 0.0036$ | 0.97<br>$\pm 0.26$ | 2029.56<br>$\pm 268.39$ | 2.08<br>$\pm 0.82$ | 146.65<br>$\pm 24.79$ |
| 5.0 | 0.0555<br>$\pm 0.0040$ | -0.0388<br>$\pm 0.0027$ | 0.0335<br>$\pm 0.0031$ | 1.01<br>$\pm 0.09$ | 1972.38<br>$\pm 163.54$ | 2.07<br>$\pm 0.34$ | 164.44<br>$\pm 23.37$ |

We also found significant correlations between the fitted elastic and viscoelastic parameters for the Mooney-Rivlin model with the concentration of agarose gels from 3-5%, see Fig. 3. These correlations indicate that *µ, β*_1_, and *τ*_2_ increase with increasing agarose concentration. However, *τ*_1_ and *β*_2_ decrease with increasing agarose concentration.

**Figure 3:**
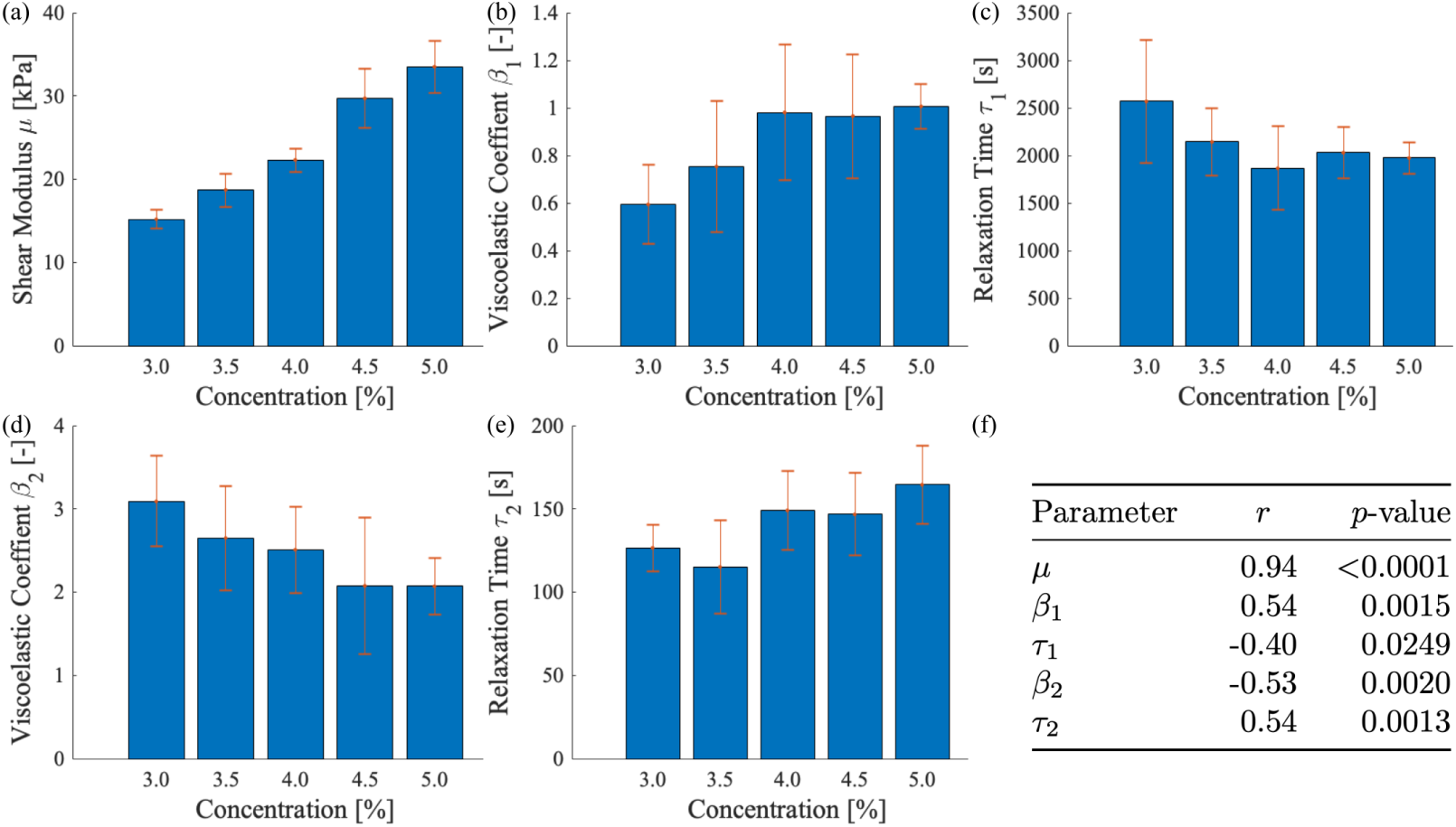
Fitted elastic and viscoelastic parameters for the Mooney-Rivlin model with the concentration of agarose gels from 3-5%: (a) shear modulus *µ*; (b) viscoelastic coefficient of long-term VP *β*_1_; (c) relaxation time of long-term VP *τ*_1_; (d) viscoelastic coefficient of short-term VP *β*_2_; (e) relaxation time of short-term VP *τ*_2_; (f) correlation *r* and significance *p* of each parameter.

### 3.2. Model Validation

Our calibrated biphasic, viscoelastic Mooney-Rivlin model successfully predicted independent experimental data from cyclic unconfined compression of specimens of agarose gels with concentrations from 3-5%, see Table 3.

**Table 3:** Comparison of FE predictions against experimental measurements from cyclic unconfined compression of specimens of agarose gels with concentrations of 3-5%, where *ϕ* is the gel concentration, *F* is the total reaction force, subscripts 0, P, and V indicate Prestrain, Peak, and Valley respectively, and subscripts FEA and EXP indicate FE predictions and experimental measurements respectively.

| $\phi$ [%] | $R^2$ [-] | $F_{0,FEA}$ [N] | $F_{0,EXP}$ [N] | $F_{P,FEA}$ [N] | $F_{P,EXP}$ [N] | $F_{V,FEA}$ [N] | $F_{V,EXP}$ [N] |
| --- | --- | --- | --- | --- | --- | --- | --- |
| 3.0 | 0.821<br>$\pm 0.153$ | 0.020 | 0.025<br>$\pm 0.008$ | 0.048 | 0.058<br>$\pm 0.010$ | -0.012 | -0.004<br>$\pm 0.010$ |
| 3.5 | 0.831<br>$\pm 0.104$ | 0.025 | 0.027<br>$\pm 0.004$ | 0.059 | 0.059<br>$\pm 0.004$ | -0.013 | -0.003<br>$\pm 0.010$ |
| 4.0 | 0.879<br>$\pm 0.074$ | 0.030 | 0.029<br>$\pm 0.008$ | 0.073 | 0.071<br>$\pm 0.010$ | -0.017 | -0.004<br>$\pm 0.004$ |
| 4.5 | 0.880<br>$\pm 0.117$ | 0.039 | 0.040<br>$\pm 0.008$ | 0.088 | 0.088<br>$\pm 0.013$ | -0.015 | -0.005<br>$\pm 0.012$ |
| 5.0 | 0.859<br>$\pm 0.080$ | 0.045 | 0.046<br>$\pm 0.006$ | 0.103 | 0.094<br>$\pm 0.010$ | -0.019 | 0.003<br>$\pm 0.008$ |

We also qualitatively compared a representative prediction against the corresponding experiment measurement, see Fig. 4(a), as well as the reaction forces at 5% prestrain, and the peaks and the valleys during the cyclic loading (predictions used mean parameters, and we compared these to the M±SD of the experiments), see Fig. 4(b).

**Figure 4:**
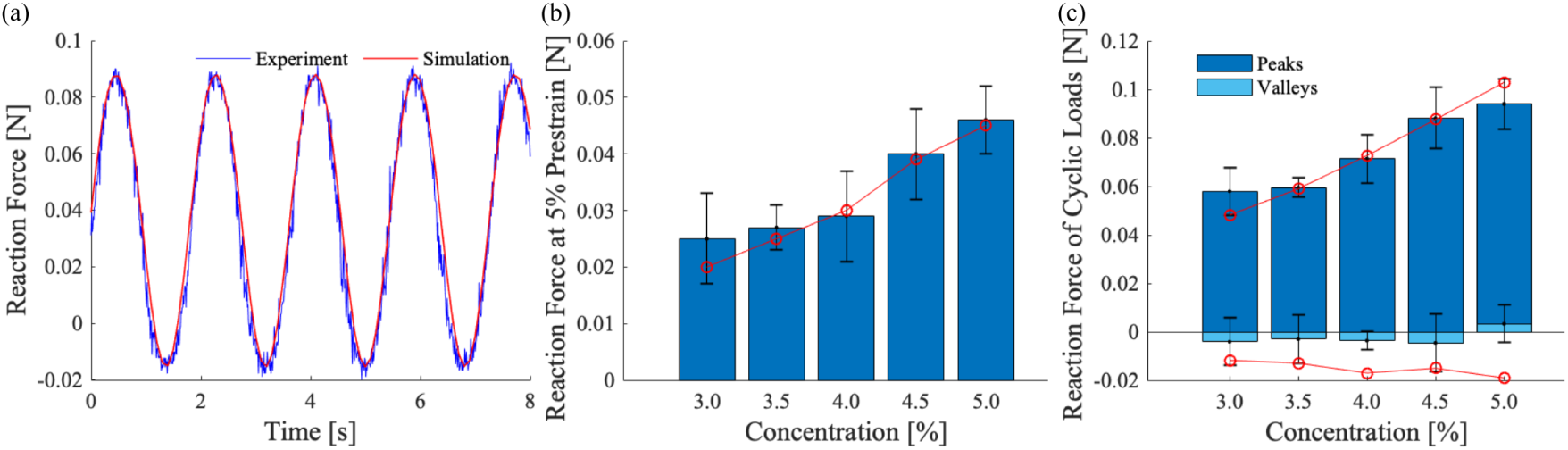
Comparison of FE predictions against experimental measurements from cyclic unconfined compression of specimens of agarose gels with concentrations of 3-5%: (a) qualitative comparison with a representative specimen; (b) comparison of the reaction force at 5% prestrain (simulation predictions in red); (c) comparison of the reaction force during the cyclic loads, peaks and valleys (simulation predictions in red.)

**Figure 5:**
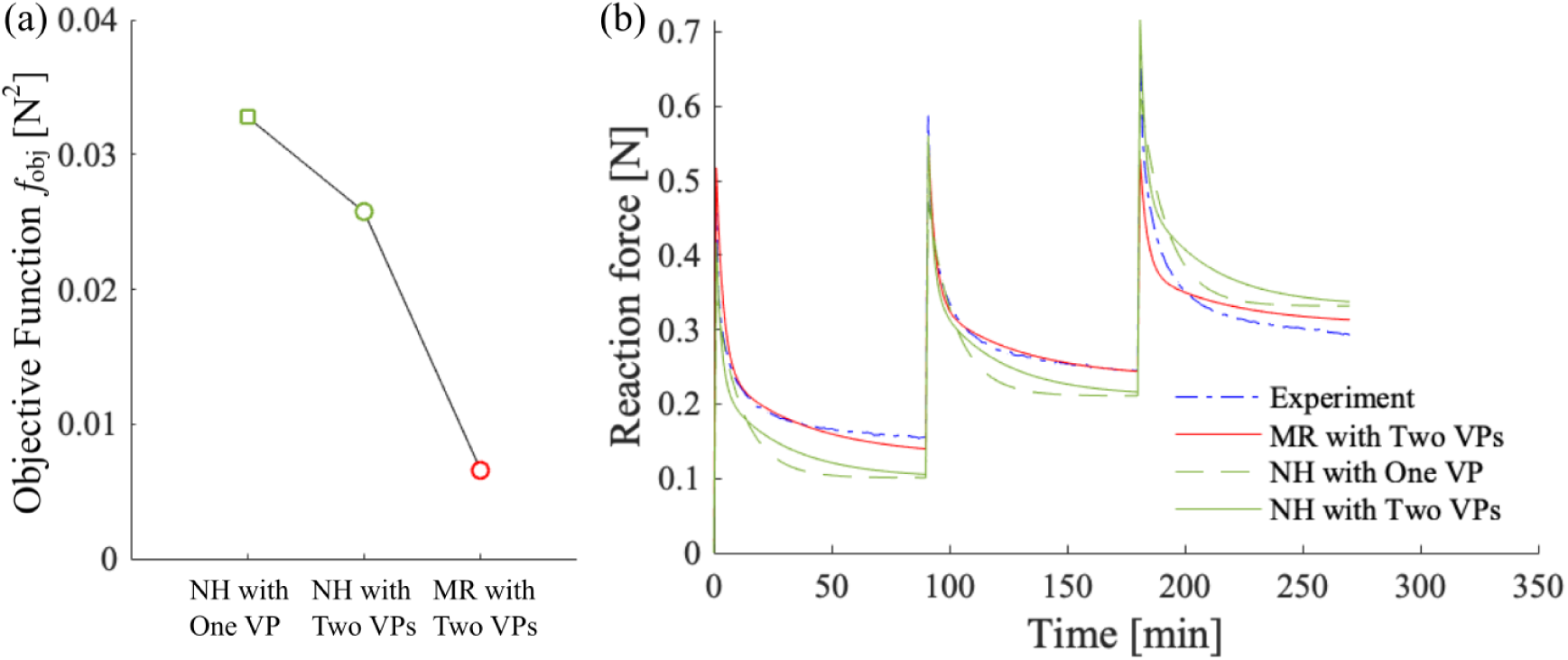
Direct comparison of the performance of the neo-Hookean and Mooney-Rivlin models: (a) values of the corresponding objective functions, (b) experimental data and predictions with fitted neo-Hookean (including one or two viscoelastic processes) and Mooney-Rivlin (including two viscoelastic processes) models for a representative agarose specimen.

## 4. Discussion

### 4.1. Model Selection and Calibration

We aimed to determine the simplest possible constitutive model which could faithfully represent our mechanical data on agarose with gel concentrations ranging from three to five percent. Prior to fitting the model using our FE-based approach we tested our FE model using an *h*-refinement test to ensure that our FE solution was sufficiently converged to represent our boundary value problem. We used a multi-step fitting process motivated by the physics of our problem. First, we fit only the elastic response within the data, i.e. the data from (near) equilibrium at the end of each stress-relaxation step. We started with a biphasic, neo-Hookean model (Mooney-Rivlin model with *C*_2_ = 0). The biphasic, neo-Hookean model stiffens with increasing compression, while our experimental data indicates that the agarose gels generally soften with increasing compression. With an additional model parameter the Mooney-Rivlin model fit the elastic portion of the experimental data much better than the neo-Hookean model, see Appendix B.

Once we fit the elastic portion of model we sought to fit the process of stress relaxation, first testing just the time-dependent poroelastic effect, then adding VPs until the model presented a good fit the experimental measurements. The poroelastic effect proved insufficient to model the stress relaxation present in the calibration data (comparison not shown). Adding one VP significantly improved the quality of the fits, while adding two VPs further improved the fits, see Fig. 2. After determining that two VPs provided the best fit we tested both one-step and two-step procedures for optimizing the parameters and improving the fits. Our subsequent analyses quantitatively demonstrated that the one-step procedure produced the best (and final) results, see Fig. 2(a).

Applying this fitting approach to all 32 specimens from a range of agarose concentrations (3-5%) we used the linear correlation coefficients to quantify trends between fitted parameters and concentrations. The shear modulus *µ* significantly increased as the concentration increased, while the other parameters did not correlate as strongly, see Figs. 3(a,f). As the concentration increased the viscoelastic coefficient of the long-term VP *β*_1_ tended to increase, see Figs. 3(b,f), while the relaxation time *τ*_1_ tended to decrease, see Figs. 3(c,f). The short-term VP showed the opposite trend, as the concentration increased the viscoelastic coefficient *β*_2_ tended to decrease, see Figs. 3(d,f), and relaxation time *τ*_2_ tended to increase, see Figs. 3(e,f).

### 4.2. Model Validation

To validate our constitutive model and model parameters we predicted results from an independent experiment including an unconfined pre-compression at 5% strain followed by cyclic compression at 3.1 to 6.9% strain. In preliminary studies we achieved a repeatable mechanical response by the third displacement-driven loading cycles. For each specimen we simulated five loading cycles and determined the peak and valley forces by averaging the response from the fourth and fifth cycles. Applying the established model to an independent boundary value problem verifies that our predictions matched the experimental data well overall with mean *R*^2^ *>* 0.82 for all concentrations of agarose, see Table 3. We also provide the comparison more intuitively in Fig. 4.

### 4.3. Limitations and Outlook

Our simulations assumed that the agarose gels remain isotropic under compression, both the solid itself and the permeation of interstitial fluid. However, agarose gels may contain fiber-like structures which may become progressively anisotropic under specific deformations like progressive uniaxial compression.

Our multi-step, physics-based optimization approach separately fits subsets of model parameters to help achieve robust convergence. Our constitutive model, fitted to experimental data on progressive stress-relaxations, was able to predict reaction forces determined from independent experiments on cyclical loading. Our fitted constitutive model has broad applications in FE modeling aimed at interpreting mechanical experiments on agarose specimens seeded with cells, cf. Jutila et al. (2015), particularly in predicting distributions of intra-gel stresses. Such experiments and FE-based analyses may allow researchers to better understand mechanobiological responses of agarose-encapsulated cells under arbitrary loads, thus elucidating the roles of stress and strain in cellular mechanotransduction.

## Acknowledgements

This material is based upon work supported by NSF CAREER 1554708, NSF CAREER 1653358, and NIH 1R01AR073964.

## Appendix A

The FE code FEBio includes an optimization scheme based on the “Levenberg-Marquardt” method for fitting constitutive models (Maas et al., 2012). The optimization scheme minimizes the objective function

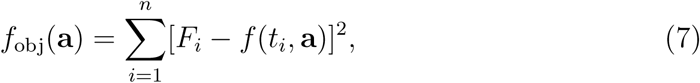

where is **a** a vector of model parameters, *n* is the number of discrete comparisons, *t*_*i*_ is a discrete time from the experiments, *F*_*i*_ is the corresponding force measured from experiments, and *f* (*t*_*i*_, **a**) is the corresponding model-predicted force (at time *t*_*i*_) using the model parameters. The optimization scheme determines the model parameters **a** that minimize the objective function *f*_obj_ by repeatedly evaluating *f* (*t*_*i*_, **a**) using FEBio to solve standard FE problems.

## Appendix B

The biphasic neo-Hookean did not fit the elastic portion of the stress-relaxation (calibration) data as well as the biphasic Mooney-Rivlin model, see Fig. 5.

